# Barcoded Bulk QTL mapping reveals highly polygenic and epistatic architecture of complex traits in yeast

**DOI:** 10.1101/2021.09.08.459513

**Authors:** Alex N. Nguyen Ba, Katherine R. Lawrence, Artur Rego-Costa, Shreyas Gopalakrishnan, Daniel Temko, Franziska Michor, Michael M. Desai

**Affiliations:** Department of Organismic and Evolutionary Biology, Harvard University, Cambridge MA 02138; NSF-Simons Center for Mathematical and Statistical Analysis of Biology, Harvard University, Cambridge MA 02138; Quantitative Biology Initiative, Harvard University, Cambridge MA 02138; Department of Physics, Massachusetts Institute of Technology, Cambridge MA 02139; Department of Molecular and Cellular Biology, Harvard University, Cambridge, MA 02138; Department of Data Science, Dana-Farber Cancer Institute, Boston MA 02215; Department of Biostatistics, Harvard T.H. Chan School of Public Health, Boston MA 02115; Department of Stem Cell and Regenerative Biology, Harvard University, Cambridge MA 02138; Center for Cancer Evolution, Dana-Farber Cancer Institute, Boston MA 02215; The Ludwig Center at Harvard, Boston MA 02215; The Broad Institute of MIT and Harvard, Cambridge MA 02139; Department of Physics, Harvard University, Cambridge, MA 02138

**Keywords:** Quantitative trait loci, polygenic traits, pleiotropy, epistasis

## Abstract

Mapping the genetic basis of complex traits is critical to uncovering the biological mechanisms that underlie disease and other phenotypes. Genome-wide association studies (GWAS) in humans and quantitative trait locus (QTL) mapping in model organisms can now explain much of the observed heritability in many traits, allowing us to predict phenotype from genotype. However, constraints on power due to statistical confounders in large GWAS and smaller sample sizes in QTL studies still limit our ability to resolve numerous small-effect variants, map them to causal genes, identify pleiotropic effects across multiple traits, and infer non-additive interactions between loci (epistasis). Here, we introduce barcoded bulk quantitative trait locus (BB-QTL) mapping, which allows us to construct, genotype, and phenotype 100,000 offspring of a budding yeast cross, two orders of magnitude larger than the previous state of the art. We use this panel to map the genetic basis of eighteen complex traits, finding that the genetic architecture of these traits involves hundreds of small-effect loci densely spaced throughout the genome, many with widespread pleiotropic effects across multiple traits. Epistasis plays a central role, with thousands of interactions that provide insight into genetic networks. By dramatically increasing sample size, BB-QTL mapping demonstrates the potential of natural variants in high-powered QTL studies to reveal the highly polygenic, pleiotropic, and epistatic architecture of complex traits.

**Significance statement:** Understanding the genetic basis of important phenotypes is a central goal of genetics. However, the highly polygenic architectures of complex traits inferred by large-scale genome-wide association studies (GWAS) in humans stand in contrast to the results of quantitative trait locus (QTL) mapping studies in model organisms. Here, we use a barcoding approach to conduct QTL mapping in budding yeast at a scale two orders of magnitude larger than the previous state of the art. The resulting increase in power reveals the polygenic nature of complex traits in yeast, and offers insight into widespread patterns of pleiotropy and epistasis. Our data and analysis methods offer opportunities for future work in systems biology, and have implications for large-scale GWAS in human populations.

## INTRODUCTION

In recent years, the sample size and statistical power of genome-wide association studies (GWAS) in humans has expanded dramatically (1–3). Studies investigating the genetic basis of important phenotypes such as height, BMI, and risk for diseases such as schizophrenia now involve sample sizes of hundreds of thousands or even millions of individuals. The corresponding increase in power has shown that these traits are very highly polygenic, with a large fraction of segregating polymorphisms (hundreds of thousands of loci) having a causal effect on phenotype (4, 5). However, the vast majority of these loci have extremely small effects, and we remain unable to explain most of the heritable variation in many of these traits (the “missing heritability” problem) (3).

In contrast to GWAS, quantitative trait locus (QTL) mapping studies in model organisms such as budding yeast tend to have much smaller sample sizes of at most a few thousand individuals (6–9). Due to their lower power, most of these studies are only able to identify relatively few loci (typically at most dozens, though see below) with a causal effect on phenotype. Despite this, these few loci explain most or all of the observed phenotypic variation in many of the traits studied (10).

The reasons for this striking discrepancy between GWAS and QTL mapping studies remain unclear. It may be that segregating variation in human populations has different properties than the between-strain polymorphisms analyzed in QTL mapping studies, or the nature of the traits being studied may be different. However, it is also possible that the discrepancy arises for more technical reasons associated with the limitations of GWAS and/or QTL mapping studies. For example, GWAS studies suffer from statistical confounders due to population structure, and the low median minor allele frequencies in these studies limit power and mapping resolution (6, 11–13). These factors make it difficult to distinguish between alternative models of genetic architecture, or to detect specific individual small-effect causal loci. Thus it may be the case that the highly polygenic architectures apparently observed in GWAS studies are at least in part artifacts introduced by these confounding factors. Alternatively, the limited power of existing QTL mapping studies in model organisms such as budding yeast (perhaps combined with the relatively high functional density of these genomes) may cause them to aggregate numerous linked small-effect causal loci into single large-effect “composite” QTL. This would allow these studies to successfully explain most of the observed phenotypic heritability in terms of an apparently small number of causal loci, even if the true architecture was in fact highly polygenic (10).

More recently, numerous studies have worked to advance the power and resolution of QTL mapping studies, and have begun to shed light on the discrepancy with GWAS (6, 14–17). One direction has been to use advanced crosses to introduce more recombination breakpoints into mapping panels (15). This improves fine-mapping resolution and under some circumstances may be able to resolve composite QTL into individual causal loci, but it does not in itself improve power to detect small-effect alleles. Another approach is to use a multiparental cross (18) or multiple individual crosses (e.g. in a round-robin mating). Several recent studies have constructed somewhat larger mapping panels with this type of design (as many as 14,000 segregants (16)); these offer the potential to gain more insight into trait architecture by surveying a broader spectrum of natural variation that could potentially contribute to phenotype. However, because multiparental crosses reduce the allele frequency of each variant (and in round-robin schemes each variant is present in only a few matings), these studies also have limited power to detect small-effect alleles. Finally, several recent studies have constructed large panels of diploid genotypes by mating smaller pools of haploid parents (e.g. a 384×104 mating leading to 18,126 F6 diploids (19)). These studies are essential to understand potential dominance effects. However, the ability to identify small-effect alleles scales only with the number of unique haploid parents rather than the number of diploid genotypes, so these studies also lack power for this purpose. Thus, previous studies have been unable to observe the polygenic regime of complex traits or to offer insight into its consequences.

Here, rather than adopting any of these more complex study designs, we sought to increase the power and resolution of QTL mapping in budding yeast simply by dramatically increasing sample size. To do so, we introduce a barcoded bulk QTL (BB-QTL) mapping approach that allows us to construct and measure phenotypes in a panel of 100,000 F1 segregants from a single yeast cross, a sample size almost two orders of magnitude larger than the current state of the art (Fig. 1A). We combined several recent technical advances to overcome the challenges of QTL mapping at the scale of 100,000 segregants: (i) unique DNA barcoding of every strain, which allows us to conduct sequencing-based bulk phenotype measurements; (ii) a highly multiplexed sequencing approach that exploits our knowledge of the parental genotypes to accurately infer the genotype of each segregant from low-coverage (<1x) sequence data; (iii) liquid handling robotics and combinatorial pooling to create, array, manipulate, and store this segregant collection in 96/384-well plates; and (iv) a highly conservative cross-validated forward search approach to confidently infer large numbers of small-effect QTL.

**Figure 1:**
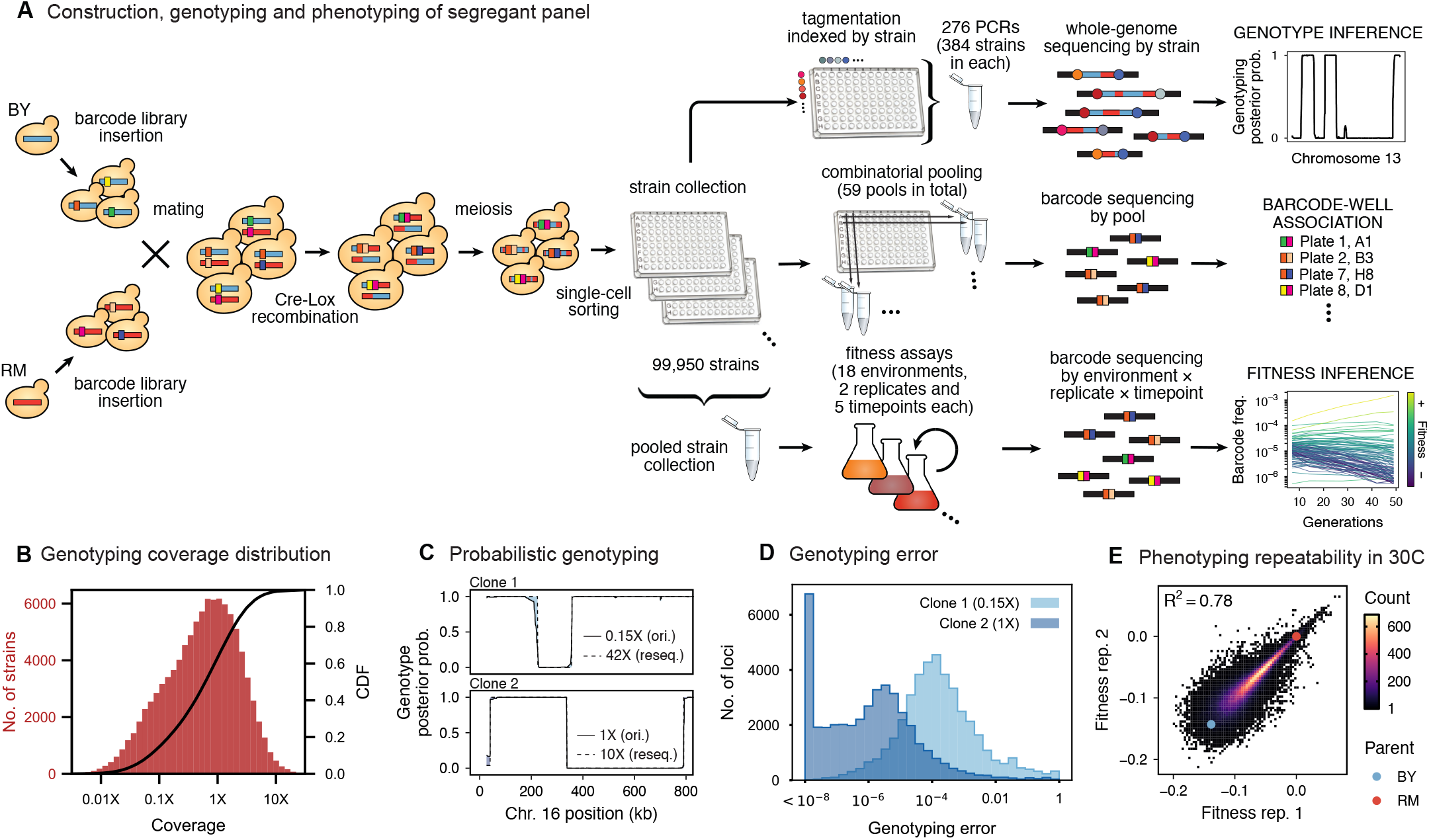
Cross design, genotyping, phenotyping, and barcode association. **A**, Construction, genotyping, and phenotyping of segregant panel. Founding strains BY (blue) and RM (red) are transformed with diverse barcode libraries (colored rectangles) and mated in bulk. Cre recombination combines barcodes onto the same chromosome. After meiosis, sporulation, and selection for barcode retention, we sort single haploid cells into 96-well plates. Top: whole-genome sequencing of segregants via multiplexed tagmentation. Middle: barcode-well association by combinatorial pooling. Bottom: bulk phenotyping by pooled competition assays and barcode frequency tracking. **B**, Histogram and cumulative distribution function (CDF) of genotyping coverage of our panel. **C**, Inferred probabilistic genotypes for two representative individuals from low coverage (solid) and high coverage (dashed) sequencing, with the genotyping error (difference between low and high coverage probabilistic genotypes) indicated by shaded blue regions. **D**, Distribution of genotyping error by SNP for the two individuals shown in (**C**). **E**, Reproducibility of phenotype measurements in 30C environment (see Fig. S2 for other environments).

Using this BB-QTL approach, we mapped the genetic basis of eighteen complex phenotypes. Despite the fact that earlier lower-powered QTL mapping studies in yeast have successfully explained most or all of the heritability of similar phenotypes with models involving only a handful of loci, we find that the increased power of our approach reveals that these traits are in fact highly polygenic, with more than a hundred causal loci contributing to almost every phenotype. We also exploit our increased power to investigate widespread patterns of pleiotropy across the eighteen phenotypes, and to analyze the role of epistatic interactions in the genetic architecture of each trait.

## RESULTS

### Construction of the barcoded segregant panel

To generate our segregant collection, we began by mating a laboratory (BY) and vineyard (RM) strain, which differ at 41,594 SNPs and vary in many relevant phenotypes (see SI). We labeled each parent strain with diverse DNA barcodes (a random sequence of 16 nucleotides), to create pools of each parent that are isogenic except for this barcode (12 and 23 pools of ∼1,000 unique barcodes in the RM and BY parental pools, respectively). Barcodes are integrated at a neutral locus containing Cre-Lox machinery for combining barcodes, similar to the “renewable” barcoding system we introduced in recent work ((20), see SI). We then created 276 sets by mating all combinations of parental pools to create heterozygous RM/BY diploids, each of which contains one barcode from each parent. After mating, we induce Cre-Lox recombination to assemble the two barcodes onto the same chromosome, creating a 32-basepair double barcode. After sporulating the diploids and selecting for doubly-barcoded haploid MATa offspring, we used single-cell sorting to select ∼100,000 random segregants and to array them into individual wells in 1,104 96-well plates. Because there are over 1 million possible barcodes per set, and only 384 offspring selected per set, this random sorting is highly unlikely to select duplicates, allowing us to produce a strain collection with one uniquely barcoded genotype in each well that can be manipulated with liquid handling robotics. Finally, we identified the barcode associated with each segregant by constructing orthogonal pools (e.g. all segregants in a given 96-well plate, all segregants in row A of any 96-well plate, all segregants from a set, etc.), and sequencing the barcode locus in each pool. This combinatorial pooling scheme allows us to infer the barcode associated with each segregant in each individual well, based on the unique set of pools in which a given barcode appears ((21); see SI).

### Inferring segregant genotypes

We next conducted whole-genome sequencing of every strain using an automated library preparation pipeline that makes use of custom indexed adapters to conduct tagmentation in 384-well plates, after which samples can be pooled for downstream library preparation (Fig. 1A). To limit sequencing costs, we infer segregant genotypes from low-coverage sequencing data (median coverage of 0.6x per segregant; Fig. 1B). We can obtain high genotyping accuracy despite such low coverage due to our cross design: because we use an F1 rather than an advanced cross, we have a high density of SNPs relative to recombination breakpoints in each individual (>700 SNPs between recombination breakpoints on average). Exploiting this fact in combination with our knowledge of the parental genotypes, we developed a Hidden Markov Model (HMM) to infer the complete segregant genotypes from this data (see SI). This HMM is similar in spirit to earlier imputation approaches (22, 23); it infers genotypes at unobserved loci (and corrects for sequencing errors and index-swapping artifacts) by assuming that each segregant consists of stretches of RM and BY loci, separated by relatively sparse recombination events. We note that this model produces probabilistic estimates of genotypes (i.e. the posterior probability that segregant genotypes is either RM or BY at each SNP; Fig. 1C), which we account for in our analysis below.

We assessed two key aspects of the performance of this sequencing approach: the confidence with which it infers genotypes, and the accuracy of the genotypes assigned. We find that at 0.1x coverage and above, our HMM approach confidently assigns genotypes at almost all loci (posterior probability of >92% of the inferred genotype at >99% of loci; see Fig. S12 and SI for a discussion of our validation of these posterior probability estimates). Loci not confidently assigned to either parental genotype largely correspond to SNPs in the immediate vicinity of breakpoints, which cannot be precisely resolved with low-coverage sequencing. To assess the accuracy of our genotyping, we conducted high-coverage sequencing of a small subset of segregants and compared the results to the inferred genotypes from our low-coverage data. We find that the genotyping is accurate (Fig. 1D), with detectable error only very near recombination breakpoints. In addition, we find that our posterior probabilities are well calibrated (e.g. 80% of the loci with an RM posterior probability of 0.8 are indeed RM; see SI). We also note that, as expected, most SNPs are present across our segregant panel at an allele frequency of 0.5 (Fig. S1), except for a few marker loci that are selected during engineering of the segregants.

### Barcoded bulk phenotype measurements

Earlier QTL mapping studies in budding yeast have typically assayed phenotypes for each segregant in their mapping panels independently, primarily by measuring colony sizes on solid agar plates (6, 14–16, 19). These colony size phenotypes can be defined on a variety of different solid media, but while they are relatively high throughput (often conducted in 384-well format), they are not readily scalable to measurements of 100,000 segregants.

Here, we exploit our barcoding system to instead measure phenotypes for all segregants in parallel, in a single bulk pooled assay for each phenotype. The basic idea is straightforward: we combine all segregants into a pool, sequence the barcode locus to measure the relative frequency of each segregant, apply some selection pressure, and then sequence again to measure how relative frequencies change. These bulk assays are easily scalable and can be applied to any phenotype that can be measured based on changes in relative strain frequencies. Because we only need to sequence the barcode region, we can sequence deeply to obtain high-resolution phenotype measurements at modest cost. In addition, we can correct sequencing errors because the set of true barcodes is known in advance from combinatorial pooling (see above). Importantly, this system allows us to track the frequency changes of each individual in the pool, assigning a phenotype to each specific segregant genotype. This stands in contrast to “bulk segregant analysis” approaches that use whole-genome sequencing of pooled segregant panels to track frequency changes of alleles rather than individual genotypes; our approach increases power and allows us to study interaction effects between loci across the genome.

Using this BB-QTL system, we investigate eighteen complex traits, defined as competitive fitness in a variety of liquid growth conditions (“environments”), including minimal, natural, and rich media with several carbon sources and a range of chemical and temperature stressors (Table 1). To measure these phenotypes, we pool all strains and track barcode frequencies through 49 generations of competition. We use a maximum likelihood model to jointly infer the relative fitness of each segregant in each assay— a value related to the instantaneous exponential rate of change in frequency of a strain during the course of the assay (Fig. 1A, lower-right inset; see SI). These measurements are consistent between replicates (average R^2^ between replicate assays of 0.77), although we note that the inherent correlation between fitness and barcode read counts means that errors are inversely correlated with fitness (Fig. 1E; Fig. S2). While genetic changes such as de novo mutations and ploidy changes can occur during bulk selection, we estimate their rates to be sufficiently low such that they impact only a small fraction of barcode lineages (see SI) and thus do not significantly bias the inference of QTL effects over the strain collection.

**Table 1:** Phenotyping growth conditions. Summary of the eighteen competitive fitness phenotypes we analyze in this study. All assays were conducted at 30°C, except when stated otherwise. YP: 1% yeast extract, 2% peptone. YPD: 1% yeast extract, 2% peptone, 2% glucose. SD: synthetic defined medium, 2% glucose. YNB: yeast nitrogen base, 2% glucose. Numbers of inferred additive QTL and epistatic interactions are also shown.

| Name | Description | Additive QTL | Epistatic QTL |
| --- | --- | --- | --- |
| 23C | YPD, 23C | 112 | 185 |
| 25C | YPD, 25C | 134 | 189 |
| 27C | YPD, 27C | 149 | 255 |
| 30C | YPD, 30C | 159 | 247 |
| 33C | YPD, 33C | 147 | 216 |
| 35C | YPD, 35C | 117 | 250 |
| 37C | YPD, 37C | 128 | 265 |
| 4NQO | SD, 0.05 g/ml 4-nitroquinoline 1-oxide | 153 | 394 |
| cu | YPD, 1mM copper(II) sulfate | 143 | 225 |
| eth | YPD, 5% (v/v) ethanol | 149 | 247 |
| gu | YPD, 6 mM guanidinium chloride | 185 | 277 |
| li | YPD, 20 mM lithium acetate | 83 | 42 |
| mann | YP, 2% (w/v) mannose | 169 | 341 |
| mol | molasses, diluted to 20% (w/v) sugars | 111 | 235 |
| raff | YP, 2% (w/v) raffinose | 167 | 221 |
| sds | YPD, 0.005% (w/v) SDS | 175 | 263 |
| suloc | YPD, 50 M suloctidil | 173 | 314 |
| ynb | YNB, w/o AAs, w/ ammonium sulfate | 145 | 303 |

### Modified stepwise cross-validated forward search approach to mapping QTL

With genotype and phenotype data for each segregant in hand, we next sought to map the locations and effects of QTL. The typical approach to inferring causal loci would be to use a forward stepwise regression (6, 22). This method proceeds by first computing a statistic such as p-value or LOD score for each SNP independently, to test for a statistical association between that SNP and the phenotype. The most-significant SNP is identified as a causal locus, and its estimated effect size is regressed out of the data. This process is then repeated iteratively to identify additional causal loci. These iterations proceed until no loci are identified with a statistic that exceeds a predetermined significance threshold, which is defined based on a desired level of genome-wide significance (e.g. based on a null expectation from permutation tests or assumptions about the numbers of true causal loci). However, although this approach is fast and simple and can identify large numbers of QTL, it is not conservative. Variables added in a stepwise approach do not follow the claimed F or chi-squared distribution, so using p-values or related statistics as a selection criterion is known to produce false positives, especially at large sample sizes or in the presence of strong linkage (24). Because our primary goal is to dissect the extent of polygenicity by resolving small-effect loci and decomposing “composite” QTL, these false positives are particularly problematic and we therefore cannot use this traditional approach.

Fortunately, due to the high statistical power of our study design, we are better positioned to address the question of polygenicity using a more conservative method with lower false discovery rate. To do so, we carried out QTL mapping through a modified stepwise regression approach, with three key differences compared to previous methods. First, we use cross-validation rather than statistical significance to terminate the model search procedure, which reduces the false positive rate. Specifically, we divide the data into training and test sets (90% and 10% of segregants respectively, chosen randomly), and add QTL iteratively in a forward stepwise regression on the training set. We terminate this process when performance on the test set declines, and use this point to define an L0-norm sparsity penalty on the number of QTL. We repeat this process for all possible divisions of the data to identify the average sparsity penalty, and then use this sparsity penalty to infer our final model from all the data (in addition, an outer loop involving a validation set is also used to assess the performance of our final model, see SI). The second key difference in our method is that we jointly re-optimize inferred effect sizes (i.e. estimated effect on fitness of having the RM versus the BY version of a QTL) and lead SNP positions (i.e. our best estimate of the actual causal SNP for each QTL) at each step. This further reduces the bias introduced by the greedy forward search procedure. Finally, the third key difference in our approach is to estimate the 95% credible interval around each lead SNP using a Bayesian method rather than LOD-drop methods, which is more suitable in polygenic architectures. We describe and validate this modified stepwise regression approach in detail in the SI. Simulations under various QTL architectures show that this approach has a low false positive rate, accurately identifies lead SNPs and credible intervals even with strong linkage, and generally calls fewer QTL than in the true model, only missing QTL of extremely small effect sizes (see SI).

### Resolving the highly polygenic architecture of complex phenotypes in yeast

We used our modified stepwise cross-validated forward search to infer the genetic basis of the eighteen phenotypes described in Table 1, assuming an additive model. We find that these phenotypes are highly polygenic: we identify well over 100 QTL spread throughout the genome for almost every trait, an order of magnitude more than that found for similar phenotypes in most earlier studies (∼0.3% of SNPs in our panel; Fig. 2, Fig. 3, Fig. S3). This increase can be directly attributed to our large sample size: inference on a downsampled dataset of 1,000 individuals detects no more than 30 QTL for any trait (see SI).

**Figure 2:**
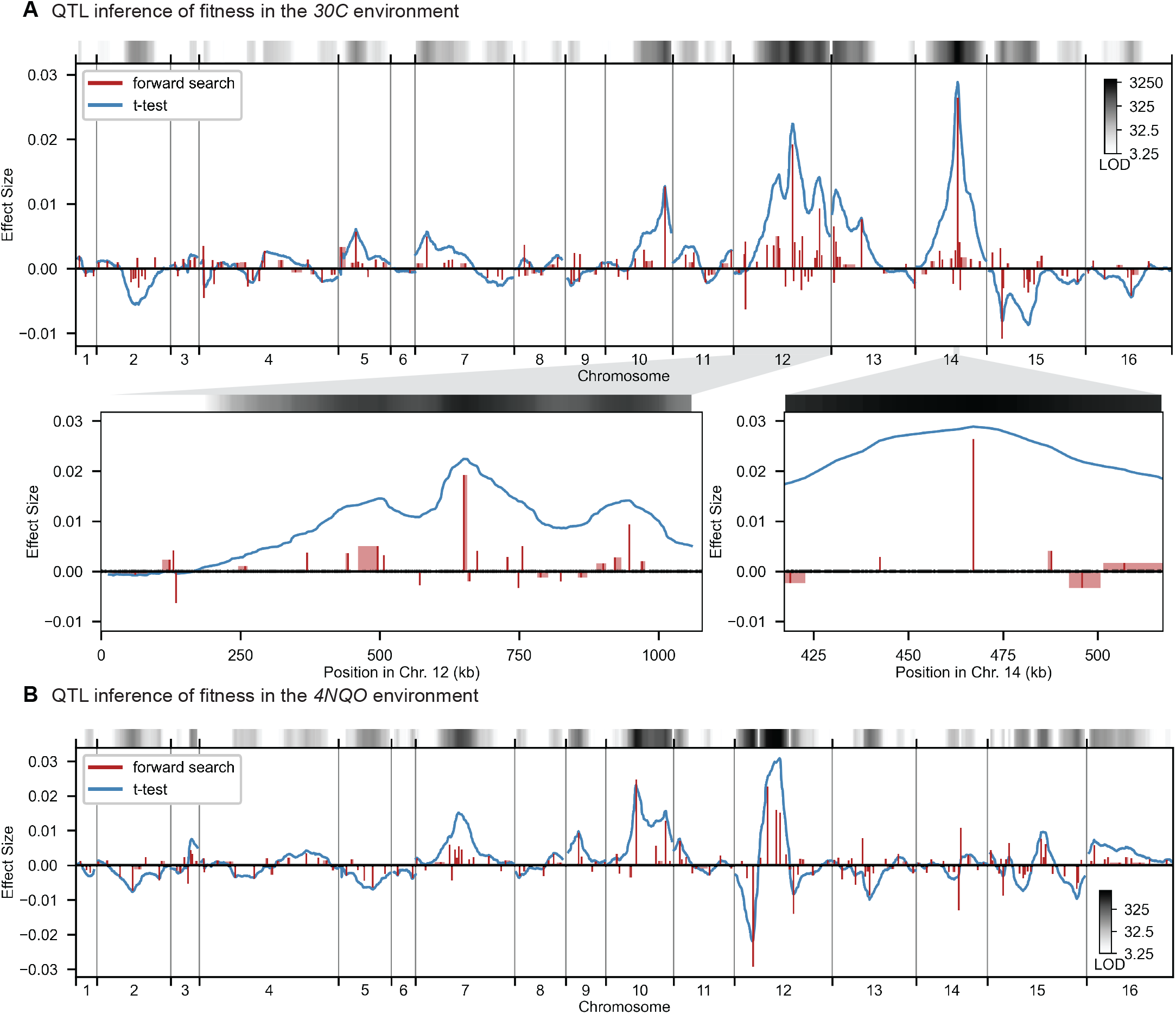
High-resolution QTL mapping. **A**,**B**, QTL mapping for **(A)** YPD at 30°C and **(B)** YPD with 4-nitroquinoline (4NQO). Inferred QTL are shown as red bars; bar height shows effect size and red shaded regions represent credible intervals. For contrast, effect sizes inferred by a Student’s t-test at each locus are shown in blue. Gray bars at top indicate loci with log-odds (LOD) scores surpassing genome-wide significance in this t-test, with shading level corresponding to log-LOD score. See Fig. S3 for other environments.

**Figure 3:**
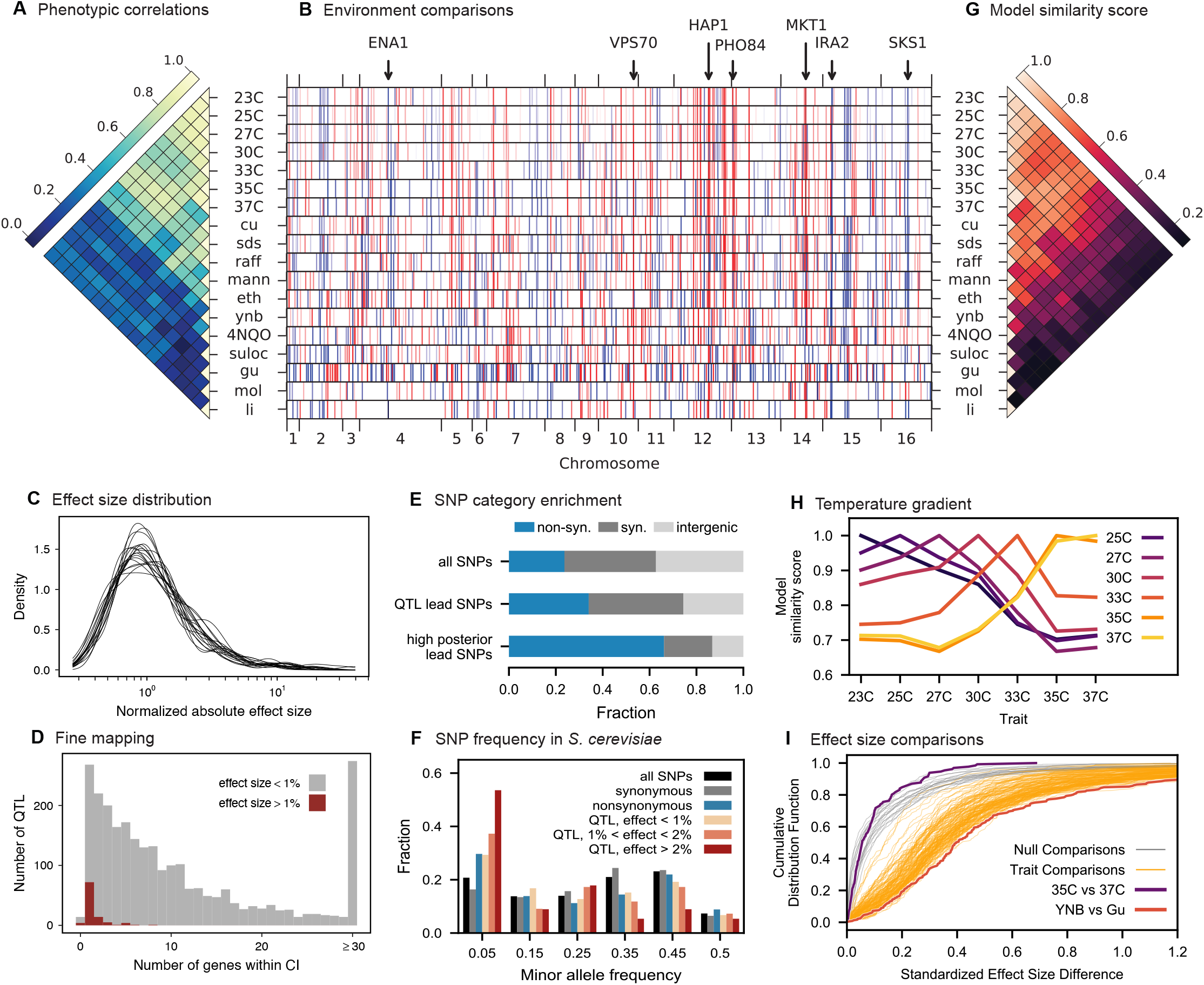
Genetic architecture and pleiotropy. **A**, Pairwise Pearson correlations between phenotype measurements, ordered by hierarchical clustering. **B**, Inferred genetic architecture for each trait. Each inferred QTL is denoted by a red or blue line for a positive or negative effect of the RM allele, respectively; color intensity denotes effect size on a log scale. Notable genes are indicated above. **C**, Smoothed distribution of absolute effect sizes for each trait, normalized by the median effect for each trait. **D**, Distribution of the number of genes within the 95% credible interval for each QTL. **E**, Distribution of SNP types. “High posterior” lead SNPs are those with >50% posterior probability. **F**, Fractions of synonymous SNPs, nonsynonymous SNPs, and QTL lead SNPs as a function of their frequency in the 1,011 Yeast Genomes panel. **G**, Pairwise model similarity scores (which quantify differences in QTL positions and effect sizes between traits; see SI) across traits. **H**, Pairwise model similarity scores for each temperature trait against all other temperature traits. **I**, Cumulative distribution functions (CDFs) of differences in effect size for each locus between each pair of traits (orange). Grey traces represent the null expectation (between cross-validation sets for the same trait). The least and most similar trait pairs are highlighted in red and purple, respectively, and indicated in the legend.

The distribution of effect sizes of detected QTL shows a large enrichment of small-effect loci, and has similar shape (though different scale) across all phenotypes (Fig. 3C), consistent with an exponential distribution above the limit of detection. This distribution suggests that further increases in sample size would reveal a further enrichment of even smaller-effect loci. While our SNP density is high relative to the recombination rate, our sample size is large enough that there are many individuals with a recombination breakpoint between any pair of neighboring SNPs (over 100 such individuals with breakpoints between each SNP on average). This allows us to precisely fine-map many of these QTL to causal genes or even nucleotides. We find that most QTL with substantial effect sizes are mapped to one or two genes, with dozens mapped to single SNPs (Fig. 3D). In many cases these genes and precise causal nucleotides are consistent with previous mapping studies (e.g. MKT1 (25), PHO84 (26), HAP1 (27)); in some cases we resolve for the first time the causal SNP within a previously identified gene (e.g. IRA2 (28)); and in others we detect novel causal genes (e.g. VPS70). However, we note that because our SNP panel does not capture all genetic variation, such as transposon insertions or small deletions, some QTL lead positions may tag linked variation rather than being causal themselves.

The SNP density in our panel and resolution of our approach highly constrain these regions of linked variation, providing guidance for future studies of specific QTL, but as a whole we find that our collection of lead SNPs displays some characteristic features of causal variants. Across all identified lead SNPs, we observe a significant enrichment of nonsynonymous substitutions, especially when considering lead SNPs with posterior probability above 0.5 (Fig. 3E; p < 10^−10^, *χ*^2^ test, df=2), as expected for causal changes in protein function. Lead SNPs are also more likely to be found within disordered regions of proteins (1.22x fold increase, p < 10^−5^), even when constrained to nonsynonymous variants (1.28x fold increase beyond the enrichment for nonsynonymous variants in disordered regions, p < 10^−4^), indicating potential causal roles in regulation (29). Lead SNP alleles, especially those with large effect size, are observed at significantly lower minor allele frequencies (MAF) in the 1,011 Yeast Genomes collection (30) compared to random SNPs (Fig. 3F; p=0.0004, Fisher’s exact test considering alleles with effect >1% and rare alleles with MAF <5%) and minor alleles are more likely to be deleterious (p=0.006, permutation test) regardless of which parental allele is rarer. These results are consistent with the view that rare, strongly deleterious alleles subject to negative selection can contribute substantially to complex trait architecture (16, 31).

### Patterns of pleiotropy

Our eighteen assay environments range widely in their similarity to each other: some groups of traits exhibit a high degree of phenotype correlation, such as rich medium at a gradient of temperatures, while other groups of traits are almost completely uncorrelated, such as molasses, rich medium with suloctidil, and rich medium with guanidinium (Fig. 3A). Because many of these phenotypes are likely to involve overlapping aspects of cellular function, we expect the inferred genetic architectures to exhibit substantial pleiotropy, where individual variants are causal for multiple traits. In addition, in highly polygenic architectures, pleiotropy across highly dissimilar traits is also expected to emerge due to properties of the interconnected cellular network. For example, SNPs in regulatory genes may affect key functional targets (some of them regulatory themselves) that directly influence a given phenotype, as well as other functional targets that may, in turn, influence other phenotypes (32).

Consistent with these expectations, we observe diverse patterns of shared QTL across traits (Fig. 3B). To examine these pleiotropic patterns at the gene level, we group QTL across traits whose lead SNP is within the same gene (or in the case of intergenic lead SNPs, the nearest gene). In total, we identify 449 such pleiotropic genes with lead SNPs affecting multiple traits (see SI). These genes encompass the majority of QTL across all phenotypes, and are highly enriched for regulatory function (Table S1) and for intrinsically disordered regions, which have been implicated in regulation (29) (p < 0.005, Fisher’s exact test, see SI). The most pleiotropic genes (Fig. S5) correspond to known variants frequently associated with quantitative variation in yeast (e.g. MKT1, HAP1, IRA2) as well as previously unidentified ones (e.g. VPS70).

The highly polygenic nature of our phenotypes highlights the difficulty in identifying modules of target genes with interpretable functions related to the measured traits (33). However, we can take advantage of our high-powered mapping approach to explore how pleiotropy leads to diverging phenotypes in different environments. Specifically, to obtain a global view of pleiotropy and characterize the shifting patterns of QTL effects across traits, we adopt a method inspired by sequence alignment strategies to match (or to leave unmatched) QTL from one trait with QTL from another trait, in a way that depends on the similarity in their effect sizes and distance between lead SNPs (see SI). From this, for each pair of environments we can find the change in effect size for each QTL, as well as an overall metric of model similarity. We find that pairwise model similarity scores recapitulate the phenotype correlation structure (Fig. 3G), including smoothly varying similarity across the temperature gradient (Fig. 3H), indicating that changes in our inferred model coefficients accurately capture patterns of pleiotropy.

For most comparisons between environments, substantial effect size changes are distributed over all QTL, indicating a broad response to the environmental shift (Fig. 3I). For example, while growth in Li (rich medium + lithium) is strongly affected by a single locus directly involved in salt tolerance (3 tandem repeats of the ENA cluster in S288C (34), corresponding to 82% of explained variance), 63 of the remaining 82 QTL are also detected in 30C (rich medium only), explaining a further 15% of variance. To some extent these 63 QTL may represent a “module” of genes with functional relevance for growth in rich medium, but their effect sizes are far less correlated than would be expected from noise or for a similar pair of environments (e.g. 30C and 27C, Fig. S6). For the temperature gradient, while we observe high correlations between similar temperatures overall, these are not due to specific subsets of genes with simple, interpretable monotonic changes in effect size. Indeed, effect size differences between temperature pairs are typically uncorrelated; thus, QTL that were more beneficial when moving from 30C to 27C may become less beneficial when moving from 27C to 25C or 25C to 23C (Fig. S6). Together, these patterns of pleiotropy reveal large numbers of regulatory variants with widespread, important, and yet somewhat unpredictable effects on diverse phenotypes, implicating a highly interconnected cellular network while obscuring potential signatures of specific functional genes or modules.

### Epistasis

To characterize the structure of this complex cellular network in more detail, numerous studies have used genetic perturbations to measure epistatic interactions between genes, which in turn shed light on functional interactions (35–40). However, the role of epistasis in GWAS and QTL mapping studies remains controversial; these studies largely focus on variance partitioning to measure the strength of epistasis, as they are underpowered to infer specific interaction terms (41). We sought to leverage the large sample size and high allele frequencies of our study to infer epistatic interactions, by extending our inference method to include potential pairwise interactions among the loci previously identified as having an additive effect (see SI). Our approach builds on the modified stepwise cross-validated search described above: after obtaining the additive model, we perform a similar iterative forward search on pairwise interactions, re-optimizing both additive and pairwise effect sizes at each step and applying a second L0-norm sparsity penalty, similarly chosen by cross-validation, to terminate the model search. We note that restricting our analysis of epistasis to loci identified as having an additive effect does not represent a major limitation. This is because a pair of loci that have a pairwise interaction but no additive effects will tend to be (incorrectly) assigned additive effects in our additive-only model, since the epistatic interaction will typically lead to background-averaged associations between each locus and the phenotype. These spurious additive effects will then tend to be reduced upon addition of the pairwise interaction term.

Using this approach, we detect widespread epistasis: hundreds of pairwise interactions for each phenotype (Fig. 4A,B, Table 1, Fig. S7), which corresponds to an average of 1.7 epistatic interactions per QTL, substantially more than has been detected in previous mapping studies (14). To interpret these epistatic interactions in the context of cellular networks, we can represent our model as a graph for each phenotype, where nodes represent genes with QTL lead SNPs and edges represent epistatic interactions between those QTL (this perspective is distinct from and complementary to Ref. (40), where nodes represent gene deletions and edges represent similar patterns of interaction). Notably, in contrast to a random graph, the epistatic graphs across phenotypes show heavy-tailed degree distributions, high clustering coefficients, and small average shortest paths (∼3 steps between any pair of genes; Fig. 4C); these features are characteristic of the small-world networks posited by the “omnigenic” view of genetic architecture (33). These results hold even when accounting for ascertainment bias (i.e. loci with large additive effects have more detected epistatic interactions; see SI).

**Figure 4:**
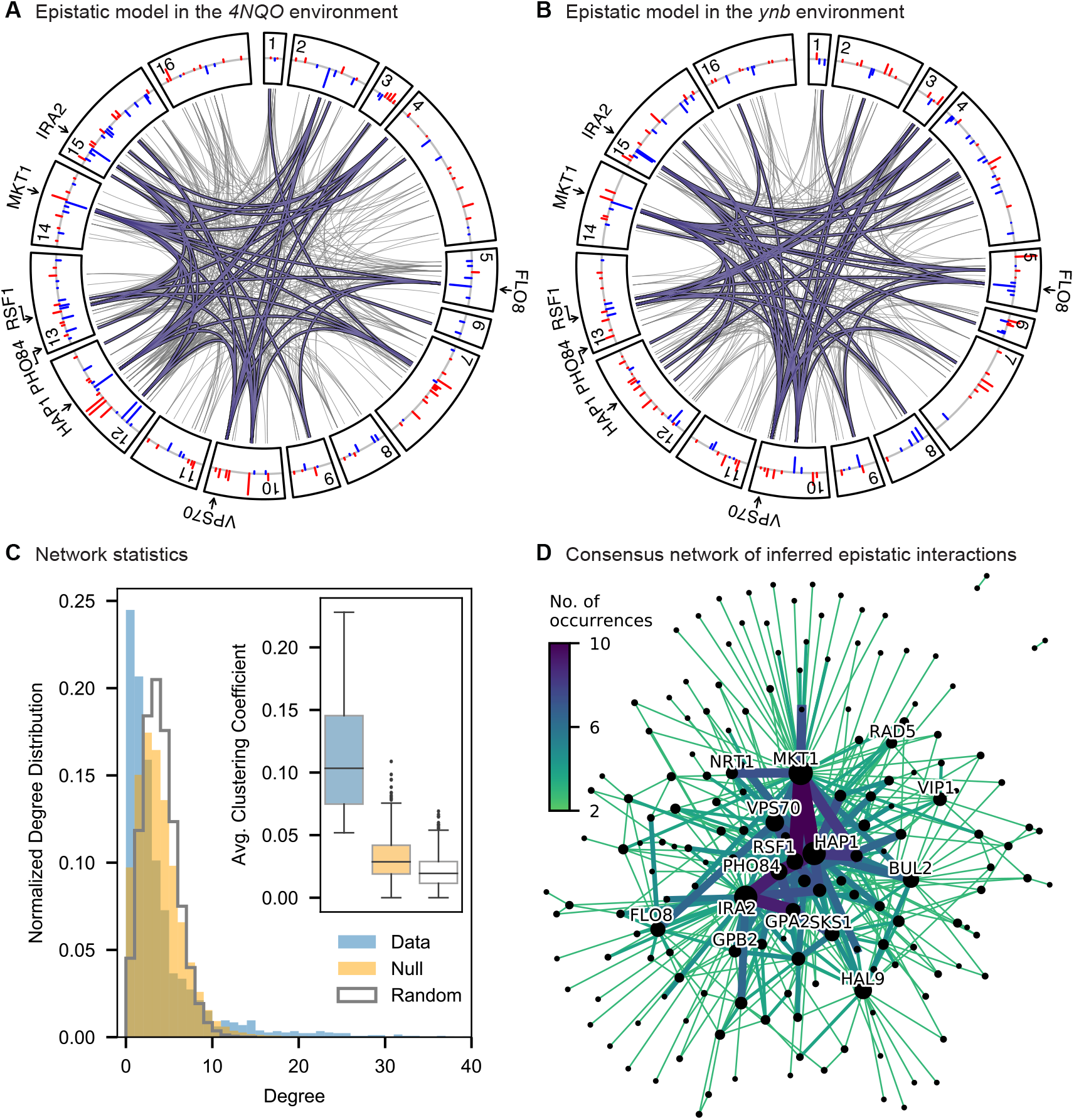
Pairwise epistasis. **A, B**, Inferred pairwise epistatic interactions between QTL (with additive effects as shown in outer ring) for (**A**) the *4NQO* environment and (**B**) the *ynb* environment. Interactions that are also observed for at least one other trait are highlighted in purple. See Fig. S7 for other environments. **C**, Network statistics across environments. The pooled degree distribution for the eighteen phenotype networks is compared with 50 network realizations generated by an Erdos-Renyi random model (white) or an effect-size-correlation-preserving null model (orange; see SI). Inset: average clustering coefficient for the eighteen phenotypes, compared to 50 realizations of the null and random models. **D**, Consensus network of inferred epistatic interactions. Nodes represent genes (with size scaled by degree) and edges represent interactions that were detected in more than one environment (with color and weight scaled by the number of occurrences). Notable genes are labeled.

We also find that hundreds of epistatic interactions are repeatedly found across environments (Fig. 4D, Fig. S8). Overall, epistatic interactions are more likely to be detected in multiple environments than expected by chance, even when considering only uncorrelated environments (p < 10^−3^, see SI), as expected if these interactions accurately represent the underlying regulatory network. Considering interactions found in all environments, we see a small but significant overlap of detected interactions with previous genome-wide deletion screens (39) (p = 0.03, *χ*^2^ = 4.46, df = 1; see SI). Taken together, these results suggest that inference of epistatic interactions in a sufficiently high-powered QTL mapping study provides a consistent and complementary method to reveal both global properties and specific features of underlying functional networks.

### Validating QTL inferences with reconstructions

We next sought to experimentally validate the specific inferred QTL and their effect sizes from our additive and additive-plus-pairwise models. To do so, we reconstructed 6 individual and 9 pairs of RM SNPs on the BY background and measured their competitive fitness in 11 of the original 18 conditions in individual competition assays (though note that for technical reasons these measurement conditions are not precisely identical to those used in the original bulk fitness assays; see SI). We find that the QTL effects inferred with the additive-only models are correlated with the phenotypes of these reconstructed genotypes, although the predicted effects are systematically larger than the measured phenotypes (Fig. 5A, cyan). To some extent, these errors may arise from differences in measurement conditions, undetected smaller-effect linked loci that bias inferred additive effect sizes, and from the confidence intervals around the lead SNP, which introduce uncertainty about the identity of the precise causal locus, among other factors. However, this limited power is also somewhat unsurprising even if our inferred lead SNPs are correct, because the effect sizes inferred from the additive-only model measure the effect of a given locus averaged across the F1 genetic backgrounds in our segregant panel. Thus, if there is significant epistasis, we expect the effect of these loci in the specific strain background chosen for the reconstruction (the BY parent in this case) to differ from the background-averaged effect inferred by BB-QTL.

**Figure 5:**
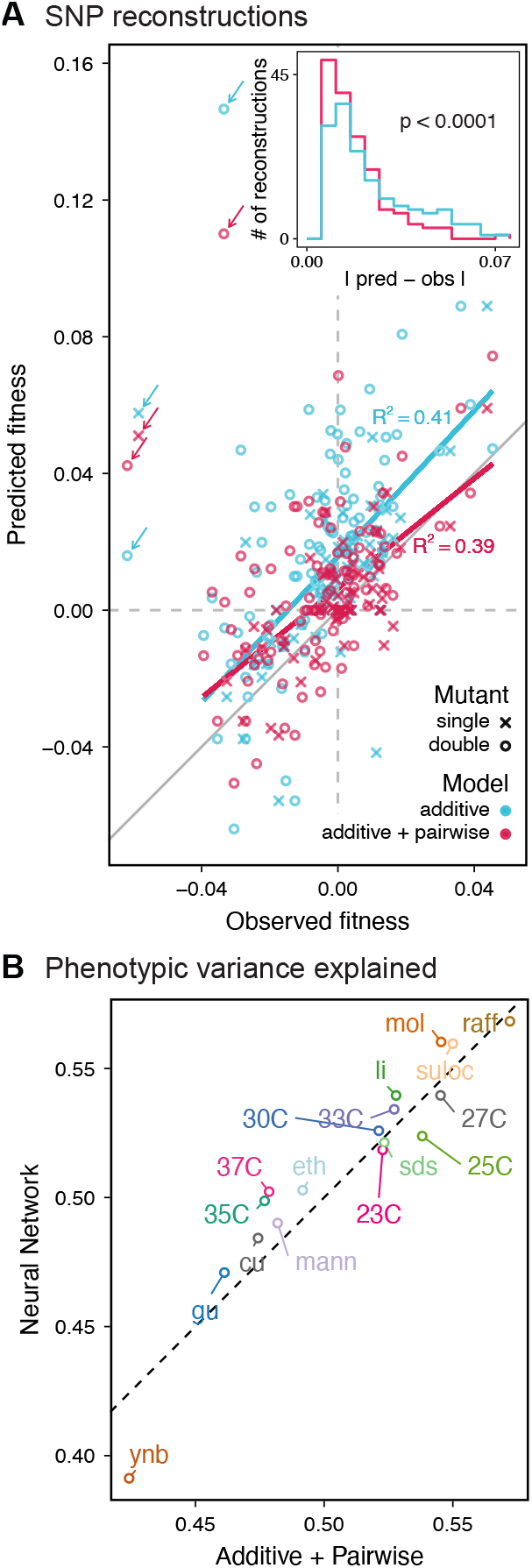
Evaluating model performance. **A**, Comparison between measured fitness effects of reconstructions of 6 single (crosses) and 9 double mutants (circles) in 11 environments, and their fitness in those environments as predicted by our inferred additive-only (cyan) or additive-plus-pairwise-epistasis models (magenta). The one-to-one line is shown in gray. R^2^ values correspond to to shown fitted linear regressions for each type of model (colored lines), excluding MKT1 mutants measured in *gu* environment (outliers indicated by arrows; see SI). Inset shows the histogram of the absolute difference between observed and predicted reconstruction fitness under our two models, with the *p*-value from the permutation test of the difference between these distributions indicated (see SI) **B**, Comparison between estimated phenotypic variance explained by the additive-plus-pairwise-epistasis model and a trained dense neural network of optimized architecture (see SI).

In agreement with this interpretation, we find that the predictions from our additive-plus-pairwise inference agree better with the measured values in our reconstructed mutants (Fig. 5A, magenta). Specifically, we find that the correlation between predicted and measured phenotypes is similar to the additive-only model, but the systematic overestimates of effect sizes are significantly reduced (Fig. 5A, inset; p < 0.0001 from permutation test; see SI). This suggests a substantial effect of nonlinear terms, although the predictive power of our additive-plus-pairwise model remains modest. As above, this limited predictive power could be a consequence of undetected linked loci or errors in the identification of interacting loci. However, it may also indicate the presence of further epistasis of higher order than the pairwise terms we infer. To explore these potential effects of higher-order interactions, we trained a dense neural network to jointly predict 17 out of our 18 phenotypes from genotype data (see SI). The network architecture involves three densely connected layers, allowing it to capture arbitrary nonlinear mappings. Indeed, we find that this neural network approach does explain slightly more phenotypic variance (on average 1% more variance than the additive-plus-pairwise QTL model, Fig. 5B, see SI), although specific interactions and causal SNPs are harder to interpret in this case. Together, these results suggest that although our ability to pinpoint precise causal loci and their effect sizes is likely limited by a variety of factors, the models with epistasis do more closely approach the correct genetic architecture despite explaining only marginally more variance than the additive model (Fig. S4), as suggested by previous studies (42).

## DISCUSSION

The BB-QTL mapping approach we have introduced in this study increases the scale at which QTL mapping can be performed in budding yeast, primarily by taking advantage of automated liquid handling techniques and barcoded phenotyping. While the initial construction of our segregant panel involved substantial brute force, this has now generated a resource that can be easily shared (particularly in pooled form) and used for similar studies aiming to investigate a variety of related questions in quantitative genetics. In addition, the approaches we have developed here provide a template for the systematic construction of additional mapping panels in future work, which would offer the opportunity to survey the properties of a broader range of natural variation. While our methods are largely specific to budding yeast, conceptually similar high-throughput automated handling systems and barcoding methods may also offer promise in studying quantitative genetics in other model organisms, though substantial effort would be required to develop appropriate techniques in each specific system.

Here, we have used our large segregant panel to investigate the genetic basis of eighteen phenotypes, defined as competitive fitness in a variety of different liquid media. The increased power of our study reveals that these traits are all highly polygenic: using a conservative cross-validation method, we find more than 100 QTL at high precision and low false-positive rate for almost every environment in a single F1 cross. Our detected QTL include many of the key genes identified in earlier studies, along with many novel loci. These QTL overall are consistent with statistical features observed in previous studies. For example, we find an enrichment in nonsynonymous variants among inferred causal loci in regulatory genes, and a tendency for rare variants (as defined by their frequency in the 1,011 Yeast Genomes collection) to have larger effect sizes.

While the QTL we detect do explain most of the narrow-sense heritability across all traits (Fig. S4), this does not represent a substantial increase relative to the heritability explained by earlier, smaller studies with far fewer QTL detected (6, 14, 16). Instead, the increased power of our approach allows us to dissect multiple causal loci within broad regions previously detected as a single “composite” QTL (Fig. 2A insets), and to detect numerous novel small-effect QTL. Thus, our results suggest that, despite their success in explaining observed phenotypic heritability, these earlier QTL mapping studies in budding yeast fail to accurately resolve the highly polygenic nature of these phenotypes. This in turn implies that the apparent discrepancy in the extent of polygenicity inferred by GWAS compared to QTL mapping studies in model organisms arises at least in part as an artifact of the limited sample sizes and power of the latter.

Our finding that increasing power reveals increasingly polygenic architectures of complex traits is broadly consistent with several other recent studies that have improved the scale and power of QTL mapping in yeast in different ways. For example, advanced crosses have helped to resolve composite QTL into multiple causal loci (15), and multiparental or round-robin crosses have identified numerous additional causal loci by more broadly surveying natural variation in yeast (16, 18). In addition, recent work has used very large panels of diploid genotypes to infer highly polygenic trait architectures, though this study involves a much more permissive approach to identifying QTL that may lead to a substantial false positive rate (19). Here, we have shown that by simply increasing sample size we can both resolve composite QTL into multiple causal loci and also identify numerous additional small-effect loci that previous studies have been underpowered to detect. The distribution of QTL effect sizes we infer is consistent with an exponential distribution up to our limit of detection, suggesting that there may be many more even smaller-effect causal loci that could be revealed by further increases in sample size.

By applying BB-QTL mapping to eighteen different fitness traits, we explored how the effects of individual loci shift across small and large environmental perturbations. Quantifying the structure of these pleiotropic effects is technically challenging, particularly for many QTL of modest effect that are not resolved to a single SNP or gene. In these cases, it is difficult to determine whether a particular region contains a single truly pleiotropic locus, or multiple linked variants that each influence a different trait. While we have used one particular approach to this problem, other choices are also possible, and ideally future work to jointly infer QTL using data from all phenotypes simultaneously could provide a more rigorous method for identifying pleiotropic loci. However, we do find that the structure of the pleiotropy in our inferred models largely recapitulates the observed correlations between phenotypes, suggesting that the causal loci we identify are largely sufficient to explain these patterns. Many of the same genes are implicated across many traits, often with similar strong effect sizes in distinct environments, and as we might expect these highly pleiotropic QTL are enriched for regulatory function. However, dividing QTL into modules that affect subsets of environments, predicting how their effect sizes change across environments (even our temperature gradient), and identifying core or peripheral genes (as in (33)) remains difficult. Future work to assay a larger number and wider range of phenotypes could potentially provide more detailed insight into the structure of relationships between traits and how they arise from shared genetic architectures.

We also leveraged the statistical power of our approach to explore the role of epistatic interactions between QTL. Previous studies have addressed this question through the lens of variance partitioning, concluding that epistasis contributes less than additive effects to predicting phenotype (14). However, it is a well-known statistical phenomenon that variance partitioning alone cannot determine the relative importance of additive, epistatic, or dominance factors in gene action or genetic architectures (41). Here, we instead explore the role of epistasis by inferring specific pairwise interaction terms and analyzing their statistical and functional patterns. We find that epistasis is widespread, with nearly twice as many interaction terms as additive QTL. The resulting interaction graphs show characteristic features of biological networks, including heavy-tailed degree distributions and small shortest paths, and we see a significant overlap with interaction network maps from previous studies despite the different sources of variation (naturally occurring SNPs versus whole-gene deletions). Notably, the set of genes with the most numerous interactions overlaps with the set of highly pleiotropic genes, which are themselves enriched for regulatory function. Together, these findings indicate that we are capturing features of underlying regulatory and functional networks, although we are far from revealing the complete picture. In particular, we expect that we fail to detect many interactions that have small effect sizes below our detection limit, that the interactions we observe are limited by our choice of phenotypes, and that higher-order interactions may also be widespread.

To validate our QTL inference, we reconstructed a small set of single and double mutations by introducing RM alleles into the BY parental background. We find that our ability to predict the effects of these putatively causal loci remains somewhat limited: the inferred effect sizes in our additive plus pairwise epistasis models have relatively modest power to predict the fitness effects of reconstructed mutations and pairs of mutations. Thus, despite the unprecedented scale and power of our study, we still cannot claim to precisely infer the true genetic architecture of complex traits. This failure presumably arises in part from limitations to our inference, which could lead to inaccuracies in effect sizes or the precise locations of causal loci. In addition, the presence of higher-order epistatic interactions (or interactions with the mitochondria) would imply that we cannot expect to accurately predict phenotypes for genotypes far outside of our F1 segregant panel, such as single- and double-SNP reconstructions on the BY genetic background. While both of these sources of error could in principle be reduced by further increases in sample size and power, it is unlikely that substantial improvements are likely to be realized in the near future.

However, despite these limitations, our BB-QTL mapping approach helps bridge the gap between well-controlled laboratory studies and high-powered, large-scale GWAS, revealing that complex trait architecture in model organisms is indeed influenced by large numbers of previously unobserved small-effect variants. We examined in detail how this architecture shifts across a spectrum of related traits, observing that while pleiotropy is common, changes in effects are largely unpredictable, even for similar traits. Further, we characterized specific epistatic interactions across traits, revealing not only their substantial contribution to phenotype but also the underlying network structure, in which a subset of genes occupy central roles. Future work in this and related systems is needed to illuminate the landscape of pleiotropy and epistasis more broadly, which will be crucial not merely for phenotype prediction but for our fundamental understanding of cellular organization and function.

## Supporting information

Supplementary Data Descriptions

## ACKNOWLEDGEMENTS

We thank the Bauer Core facility at Harvard and the Broad Institute Genomic Services sequencing core for assistance with sequencing. We thank Alan Moses, Andrew Murray, Hunter Fraser, Dan Rice, and members of the Desai lab for comments on the manuscript. A.N.N.B. acknowledges support from the National Science and Engineering Research Council of Canada (NSERC). K.R.L. acknowledges support from the Fannie & John Hertz Foundation Graduate Fellowship Award, the National Science Foundation (NSF) Graduate Research Fellowship Program, and fellowship award #1764269 from the NSF-Simons Center for Mathematical and Statistical Analysis of Biology at Harvard. M.M.D. acknowledges support from grant PHY-1914916 from the NSF and grant GM104239 from the National Institute of Health (NIH). The computations in this paper were run on the Faculty of Arts and Sciences Research Computing (FASRC) Cannon cluster supported by the FAS Division of Science Research Computing Group at Harvard University.

## AUTHOR CONTRIBUTIONS

A.N.N.B., K.R.L., A.R.-C., and M.M.D. designed the project; A.N.N.B., K.R.L., and A.R.-C. constructed and genotyped the panel; A.N.N.B., K.R.L., A.R.-C, and S.G. conducted phenotyping; A.N.N.B., K.R.L., and A.R.-C. developed inference methods and analyzed the data; D.T. and F.M. developed the neural-network model; A.N.N.B., K.R.L., A.R.-C., and M.M.D. wrote the paper.

## AUTHOR INFORMATION

The authors declare no competing financial interests.

## DATA AVAILABILITY

Raw sequencing reads will be uploaded to NCBI SRA upon acceptance. Inferred genotype and phenotype data (25 GB) are publicly available at https://utoronto-my.sharepoint.com/:f:/g/personal/alex_nguyenba_utoronto_ca/ElIGeEeuZo9Ok9d03NPm-QAB-ErMYC-stQD14Vd_QKlUGg?e=cYuBKG. Inferred QTL and other data are compiled in Supplementary Data Files S1-8.

## CODE AVAILABILITY

Custom code and freely available software were used to conduct analyses. All software packages and algorithms used are fully described in the Supplementary Information, and code will be made available upon request.

