## Supplementary Data Descriptions for "Barcoded Bulk QTL mapping reveals highly polygenic and epistatic architecture of complex traits in yeast"

**Supplementary Data Files**

**S1: Inferred additive QTL.** For each trait, a list of all inferred additive QTL is ranked in order of decreasing effect size magnitude. For each QTL, we provide the effect size; SNP list index of the lead SNP, credible interval (CI) start, and CI end; chromosome; chromosome position (in basepairs) of the lead SNP, CI start, and CI end; and the list of genes with coding regions at least partially overlapping the CI (gene names are given when possible, otherwise ORF names are given).

**S2: Pleiotropic genes.** A list of genes containing a lead SNP in two or more traits (“pleiotropic genes”), ranked by the number of traits. For each gene, we give the number of traits in which a lead SNP was detected; the chromosome; the gene name; the number of consensus genes (genes which overlap the intersection of credible intervals from all traits); the names of consensus genes; the lead SNP index identified in each of the traits (blank entries for traits in which the gene was not detected); the lead SNP chromosome position in basepairs in each of the traits; and the effect size in each of the traits. Following the list of pleiotropic genes, we provide a list of remaining QTL detected in only one environment, with associated data in the same format.

**S3: Inferred epistatic QTL.** For each trait, separate lists of the additive QTL and pairwise epistatic QTL inferred in additive-plus-pairwise models, ranked in order of decreasing effect size magnitude. For additive QTL, we provide the effect size; SNP list index of the lead SNP; chromosome; and gene in which the lead SNP is located (uppercase gene/ORF names indicate SNPs in coding regions, and lowercase gene/ORF names indicate that the SNP is intergenic but located closest to the gene given). For epistatic QTL, we provide the effect size; SNP list indices of both lead SNPs; chromosomes of both lead SNPs; chromosome positions in basepairs of both lead SNPs; and the gene name of both lead SNPs (formatted as for additive QTL). For both additive and epistatic QTL, we denote QTL located at selection markers (or immediately neighboring genes) in grey and place them at the bottom of the list; see SI.

**S4: Multiplicity of epistatic QTL across traits**. A list of gene pairs that are observed in epistatic interactions across all traits, ranked by the number of traits in which they occur (“edge multiplicity”). For each gene pair, we provide the gene names, corresponding to those in file S3; edge multiplicity (number of traits in which this pair of genes had a detected interaction); node pair multiplicity (number of traits in which this pair of genes were both detected as additive QTL, whether or not an interaction was detected); and the list of traits in which an interaction was detected. QTL located at selection markers (or immediately neighboring genes) are denoted in grey and placed at the bottom of the list; see SI.

**S5: GO terms of pleiotropic genes.** Full results of GO analysis on pleiotropic genes (see SI).

**S6: Variance partitioning.** Variance partitioning, including error, additive genetic, epistatic genetic components, and neural-network on both raw and resampled phenotype data. See SI for full definitions and discussion of all components.

**S7: Variant reconstructions.** Measured and predicted effects for the 6 SNPs reconstructed in the BY background, for the 11 traits assayed.

**S8: SNP list.** Our final list of 41,594 SNPs. We provide the chromosome; the SNP index (note these are indexed from 1, while SNP indices in other files are indexed from 0); the chromosome position in basepairs; and the BY and RM alleles.
